# Patterning of the vertebrate head in time and space by BMP signalling

**DOI:** 10.1101/592451

**Authors:** Kongju Zhu, Herman P. Spaink, Antony J. Durston

## Abstract

How head patterning is regulated in vertebrates is yet to be understood. In this study, we show that frog embryos injected with Noggin at different blastula and gastrula stages had their head development sequentially arrested at different positions. When timed BMP inhibition was applied to BMP-overexpressing embryos, the expression of five genes: *xcg-1* (a marker of the cement gland, which is the front-most structure in the frog embryo), *six3* (a forebrain marker), *otx2* (a forebrain and mid-brain marker), *gbx2* (an anterior hindbrain marker) and *hoxd1* (a posterior hindbrain marker) were sequentially fixed. These results suggest that timed interactions between BMP and anti-BMP are involved in patterning the vertebrate head progressively in time and space. Since the above genes are not expressed sequentially, there may be a BMP dependent gene sequence during head patterning that can be arrested by BMP inhibition and regulate the specification of positional values in the head.

## Introduction

During early development, the vertebrate embryo is patterned from anterior to posterior in a temporally progressive manner (Nieuwkoop 1952; Eyal-Giladi 1954; Gamse & Sive 2000; Gamse & Sive 2001; Stern *et al*. 2006): anterior tissues are specified early, and more posterior tissues are determined progressively later. Whereas coordination between temporal and spatial control of anterior-posterior (A-P) patterning is evident, a thorough understanding of the underlying mechanisms is still lacking in vertebrates.

In frog, there is evidence for a BMP/anti-BMP dependent time-space translation mechanism that regulates trunk-tail patterning by *Hox* genes (Wacker *et al*. 2004a; Durston & Zhu 2015). In this mechanism, *Hox* genes are sequentially activated in a high BMP region of the mesoderm (non-organiser mesoderm) (Wacker *et al*. 2004b), where their expression is dynamic and unstable. As the mesoderm involutes during gastrulation, *Hox* expressing cells are successively exposed to signals from the Spemann organiser, resulting in the *Hox* sequence being fixed at different points along the forming axis. In this way, the timing information encoded by *Hox* genes is translated into a spatial pattern. The putative organiser signals that stabilise *Hox* codes are BMP antagonists, e.g. Noggin (Smith & Harland 1992) and Chordin (Sasai *et al*. 1994), because these mimic the function of the organiser in dorsalising the embryo (Smith *et al*. 1993; Khokha *et al*. 2005), inducing secondary axes (Spemann & Mangold 1924; Sasai *et al*. 1994; Fang *et al*. 2000), and rescuing A-P axes in ventralised embryos (Sasai *et al*. 1994; Wacker *et al*. 2004a). Notably, timed Noggin treatments in ventralised embryos not only rescue the A-P axis, but also the spatial pattern of *Hox* gene expression (Wacker *et al*. 2004a). This conclusion is further supported by a recent study in chick, which reported the fixation of *Hox* codes in explanted posterior primitive streak (containing high BMP mesoderm) by Noggin treatments (Dias *et al*. 2014). Together, these findings suggest that BMP signalling is involved in patterning the trunk-tail part of the axis by (directly or indirectly) regulating *Hox* gene expression.

In the rescue experiments mentioned above (Sasai *et al*. 1994; Wacker *et al*. 2004a), Noggin and Chordin treatments can also rescue the head part of the axis, suggesting that BMP signalling may also be involved in patterning the head. Using heat-shock inducible *chordin* transgene lines (Tg (*hsp70:chd*)), Hashiguchi et al. have shown in zebrafish that the expression of *six3* (a forebrain marker) (Kobayashi *et al*. 1998), *otx2* (a forebrain and mid-brain marker) (Li *et al*. 1994; Mori *et al*. 1994), *gbx1* (the counterpart of Xenopus *gbx2*; a rostral hindbrain marker) (Rhinn *et al*. 2003), and *hoxb1b* (a caudal hindbrain marker) (Alexandre *et al*. 1996) are sequentially induced by timed anti-BMP treatment from mid-blastula to early gastrula stages (Hashiguchi & Mullins 2013). This is consistent with the observations that timed Noggin injections in ventralised embryos rescued different portions of the A-P axis in frog (Wacker *et al*. 2004a), and that progressively later anti-BMP (Chordin) treatments resulted in progressively more posterior axis defects in zebrafish (Tucker *et al*. 2008). These findings raise an interesting question: is BMP signalling involved in progressively patterning the head (brain) region of the embryo?

In deuterostomes, the front-most portion of the A-P axis is not the head, but the extreme anterior domain (EAD), a region wherein ectoderm and endoderm directly juxtapose in the early embryo (Jacox *et al*. 2014). In frog, this region gives rise to three organs, the cement gland (CG), the primary mouth, and the anterior pituitary gland (Dickinson & Sive 2007). Among them, the cement gland is an ectodermal organ that lies anterior to any neural tissue (Sive *et al*. 1989). The formation of CG can be affected by perturbations of the development of dorsal mesoderm (the Spemann organiser) (Scharf & Gerhart 1983; Kao & Elinson 1988), suggesting a requirement for organiser signals in the formation of this anterior-most structure. It would therefore be interesting to see if CG formation is also regulated by BMP signalling.

To test the role of BMP signalling in head patterning, we did timed anti-BMP treatments in both wild-type (WT) and ventralised frog embryos. This resulted in sequential arrest (in WT embryos) or rescue (in ventralised embryos) of head and EAD patterning at different values, suggesting that a timing mechanism, which is BMP dependent and can be converted into spatial patterns by anti-BMP signals, may be involved in patterning the vertebrate head-EAD.

## Results and discussion

### Timed anti-BMP treatment arrests head patterning at different positions

To examine the role of BMP signalling in head formation, we injected Noggin protein, an antagonist of BMP (Smith & Harland 1992; Zimmerman *et al*. 1996), into the blastocoel of the embryo at stage 8, 9, 10, 10.5 and 11 (from blastula to gastrula stage) (Fig 1). Embryos injected with Noggin at stage 8 formed a ball of tissue with a large cement gland. Embryos injected at stage 9 also showed a blob of tissue, but the cement gland was much smaller. When Noggin was injected at st.10, a visible, short head (half head) was formed in the embryo. Injection of Noggin at st.10.5 and 11 resulted in the formation of more posterior structures, truncating head formation at the hindbrain and neck region, respectively. Therefore, sequentially later Noggin treatments arrested head and EAD formation at more and more posterior positions, suggesting that the EAD and head are patterned gradually in a timed fashion. In zebrafish, sequential BMP inhibition results in axial defects at progressively more posterior positions: later anti-BMP treatment affects posterior positions but not anterior positions (Tucker *et al*. 2008). Together, these results suggest that the process involved in head-tail patterning can be stopped sequentially by timed BMP inhibition.

**Fig 1.**
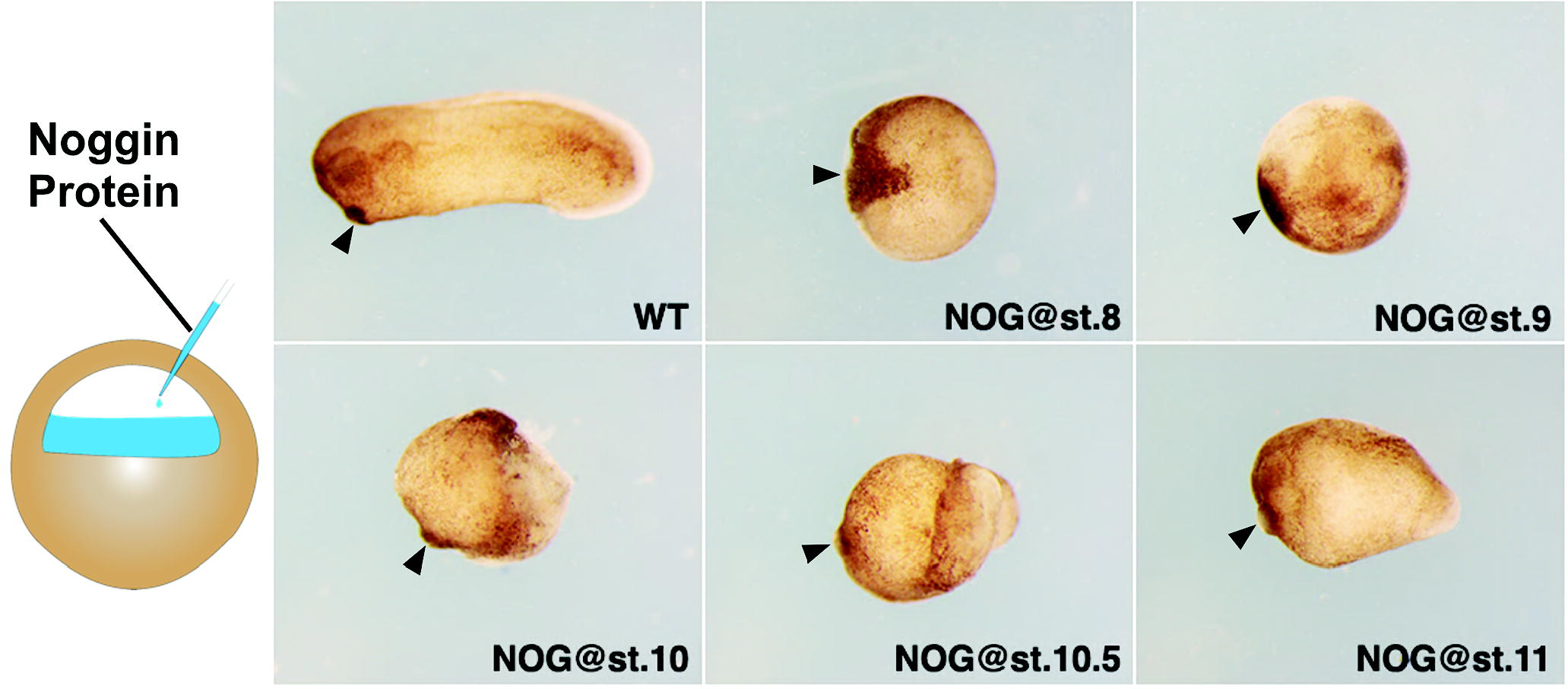
Timed Noggin-injection in WE embryos resulted in progressive arrest of head patterning. 200nl 1ng/µL noggin was injected into the blastocoel of the embryo at different stages. Anterior is to the left and dorsal is up. Black arrows point to the position of the cement gland.

### Timed anti-BMP treatment in ventralised embryos rescued different portions of the head

The above observations led us to postulate that the “head timer” is BMP-dependent and can be sequentially fixed by BMP inhibition, resulting in positional values being sequentially specified. To test this, we did timed anti-BMP treatment in ventralised frog embryos (high BMP) (Fig. 2).

**Fig 2.**
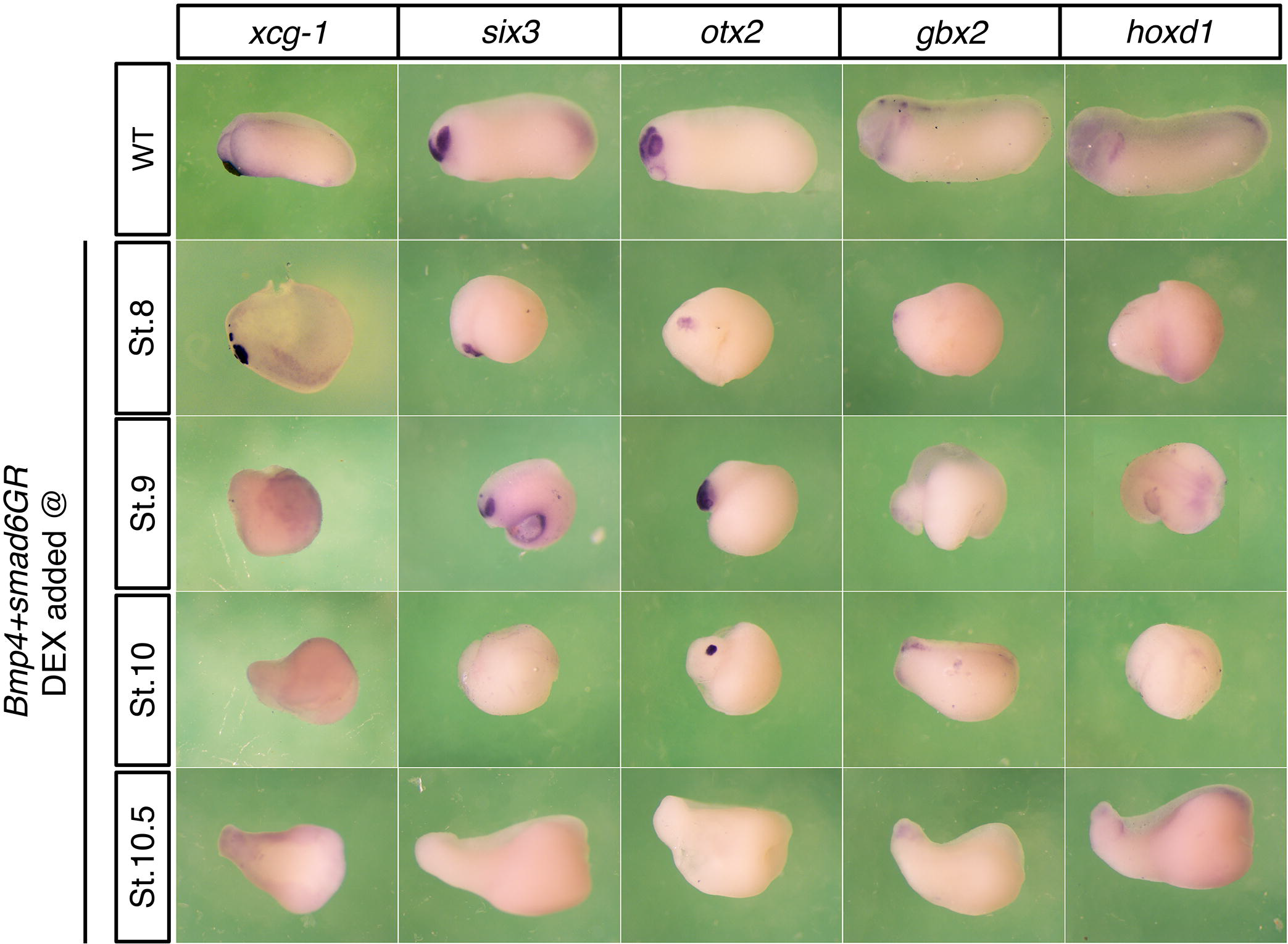
Timed anti-BMP treatment in ventralised embryos led to sequential fixation of anterior genes. The expression of *xcg-1, six3, otx2, gbx2*, and *hoxd1* in *bmp4*-injected embryos that were subjected to Smad6 treatment at different stages.

In *Xenopus*, ventralisation can be achieved by *BMP* overexpression (Dale *et al*. 1992; Jones *et al*. 1992; Clement *et al*. 1995; Schmidt *et al*. 1995), while dorsalisation can be achieved by BMP inhibition (Smith *et al*. 1993; Zhu *et al*. 2017). In our experiments, injection of the frog embryo with 2ng *bmp4* resulted in complete ventralisation, showing a blob of tissue that had no axis (Fig S1). When the embryo was injected with the same amount of *smad6*, an inhibitory Smad that can interfere with BMP pathway (Imamura *et al*. 1997; Hata *et al*. 1998; Goto *et al*. 2007), however, it displayed a dorsalised phenotype (Fig S1). We then did anti-BMP treatments in BMP4-ventralised embryos at different stages using a Smad6GR construct (Marom *et al*. 2005). Timed activation of this GR construct with dexamethazone induced timed BMP pathway inhibition by *smad6* activation. Timed Smad6 treatment fixed five anterior markers sequentially: it strongly fixed *xcg-1* at stage 8, *six3* at stage 8 and 9, *otx2* at stage 9 and 10, *gbx2* at stage 10 and 10.5, and *hoxd1* at stage 10.5 (Fig. 3). The sequential fixation of these genes fits well with the timing observed in Fig 1, suggesting that the head-EAD is progressively patterned by timed BMP/anti-BMP actions.

**Fig 3.**
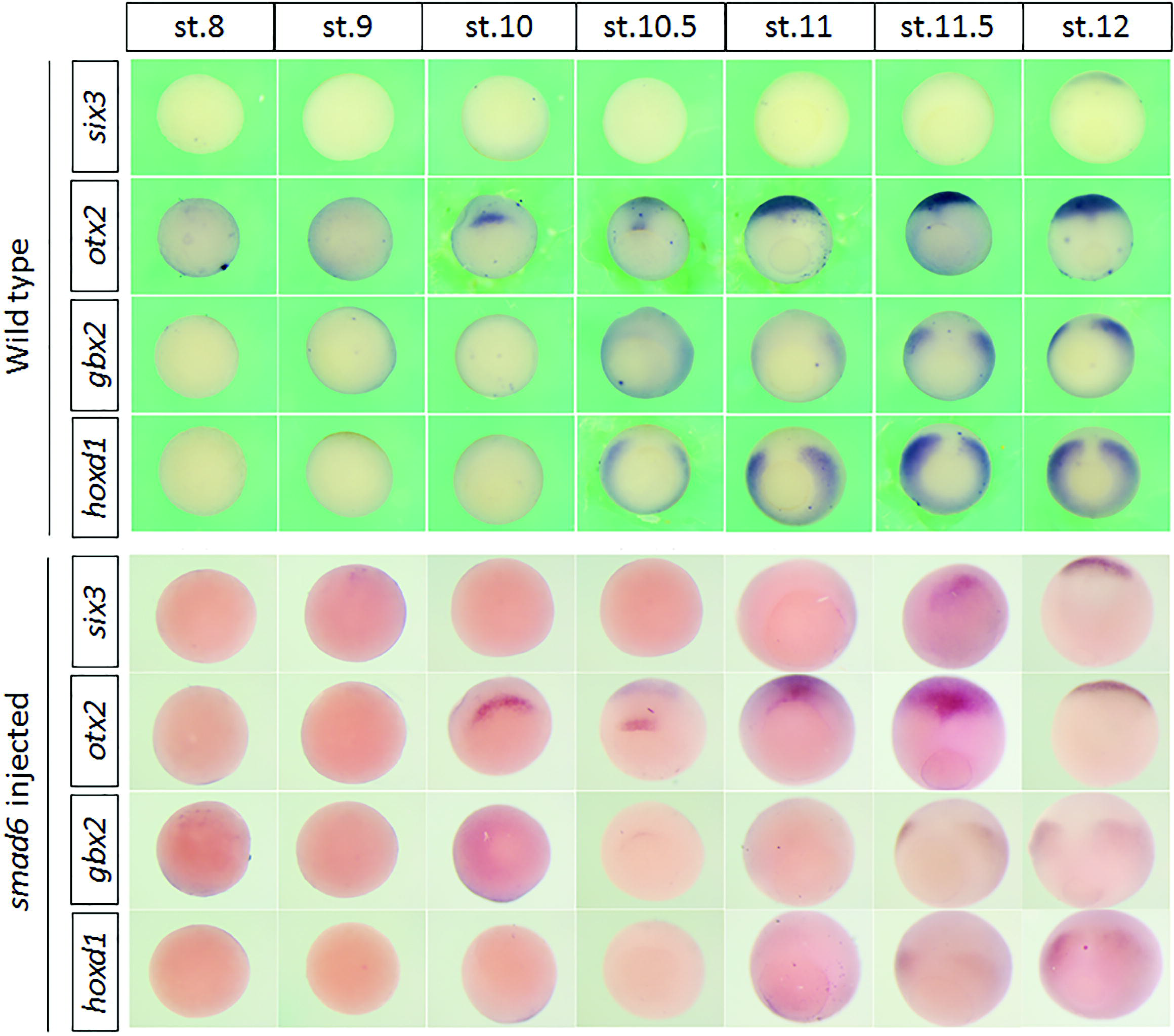
The expression of *six3, otx2, gbx2* and *hoxd1* at different stages in wild-type and *smad6*-injected embryos. Embryos are vegetal views with dorsal to the top, smad6 injected at 2-cell or 4-cell stage.

It is worth noting that, both in Hashiguchi’s study and in this study, the last and most posterior component of the head gene sequence is *hox1*: *hoxb1* and *hoxd1*, respectively. Since *hox1* is the earliest, most anterior component of a previously elucidated *Hox* time-space sequence, the spatial arrangement of these early induced head genes is clearly complementary to and continuous with the later, more posterior *Hox* gene sequence. It is also clearly continuous with a yet earlier, more anterior EAD sequence. In short, a BMP*-*anti BMP time-space sequence covers the whole A–P axis.

### The timing of A-P markers is disrupted in *smad6*-injected embryos

During trunk patterning, collinearity causes *Hox* genes to be expressed in a 3’ to 5’ order, and that more 3’ genes are expressed earlier and more anteriorly than/to more 5’ ones (Lewis 1978; Duboule & Dolle 1989; Graham *et al*. 1989). The temporally collinear expression of *Hox* genes has been proposed to serve as a timer, which can be interpreted and translated into spatial information (Wacker *et al*. 2004a). Since the anterior genes are sequentially fixed earlier than *Hox* genes by anti-BMP treatment, it is natural to think that these genes are also expressed in a temporal sequence, which could complement the *Hox* sequence to constitute an integrative timer. In addition, the very early induction of *xcg1*, Fig. 2 indicates that the EAD is timed very early in the same sequence as the head. However, although these genes showed a spatial sequence of expression along the A-P axis (Fig S2), the endogenous timed activation of these genes did not strictly correspond to their spatial order along the A-P axis (Fig 3). For example, *six-3* demarcates the most anterior border of the developing neural plate (Oliver *et al*. 1995), but it was expressed at the end of gastrulation, much later than the other genes. The expression domain of *gbx2* is anterior to that of *hoxd1*, but it was also not expressed earlier than *hoxd1*. Moreover, unlike Hox genes, which are expressed in ventral and lateral mesoderm during gastrulation, the earliest expression of *six3* and *otx2* was located at the dorsal side of the embryo. The expression kinetics and expression locations of these genes therefore make them less likely to be “timer genes” or “fixation genes” themselves (at least not all of them are). They are presumably downstream of the timer. Even so, however, their timing was disrupted by *smad6* injection (injection at 2-cell or 4-cell stage, Smd6 remaining available till much later) (Fig 3). For example, the expression of *six3* in *smad6*-injected embryos was advanced to stage 11.5 from stage 12. Expression of *gbx2* and *hoxd1* was significantly reduced and only detectable from stage 11.5, whereas their expression in WT embryos was observed earlier (at stage 10.5). These results are unsurprising. The anterior *six3* would be induced (evidently prematurely) by this early BMP inhibition. *gbx2* and *hoxd1* would be inhibited and are evidently delayed because the axial time and position has been fixed at early/anterior by this early anti-BMP treatment. Although these anterior genes may only serve as positional markers, the time of their expression is also of crucial importance. For example, *gbx2* shows a significant effect on head development when ectopically expressed at stage 9 and 10. The effect gets less drastic when it is expressed at later stages, e.g. stage 12 and 13 (Tour *et al*. 2002). Therefore, these results further emphasise the importance of BMP signalling in regulating head patterning.

### Hypothesis for head patterning by BMP signalling

The results in this study suggest two aspects of head patterning. First, the vertebrate head and EAD are patterned in a temporally progressive manner: the EAD is patterned first, followed by the patterning of the forebrain, the midbrain, the hindbrain and the neck. Second, BMP signalling is involved in patterning the head in time and space, which can be seen from the progressive arrest of head and EAD formation by timed anti-BMP treatments (Fig 1) and from sequential fixation of anterior marker genes (*xcg-1, six3, otx2, gbx2*, and *hoxd1*) by anti-BMP (Fig 2). However, the molecular mechanism, by which BMP signalling regulates head-EAD patterning, is not yet clear. It is likely a similar mechanism to that which patterns the trunk-tail part of the axis is involved (Wacker *et al*. 2004a; Durston & Zhu 2015) (Fig 4). A key component of this mechanism is a BMP dependent timer, which is temporal gene sequence. The existence of such a timer has been demonstrated in gastrula NOM mesoderm for the Xenopus *Hox* sequence. During trunk-tail patterning, the timer is likely to be regulated by *Hox* genes (Wacker *et al*. 2004a; Durston & Zhu 2015). The timer genes involved in head-EAD patterning and the location of the timer, however, are so far unknown. Some of the anterior head makers (e.g. *six3, gbx2*) that we examined in our study presumably do not regulate timing or fixation because they were not sequentially expressed endogenously (Fig 3). Since these genes could be fixed sequentially by anti-BMP treatments, a possible explanation is that sequential states of the currently unknown “head timer” are fixed sequentially by anti-BMP signals and these then regulate the expression of anterior marker genes and hence progressive patterning of the head-EAD. In conclusion, the results in study argue for a vital role for BMP/anti-BMP in patterning the vertebrate head in time and space. We postulate that a ventral BMP dependent timer sequentially exposes and dorsal anti BMP sequentially fixes an early-anterior to late-posterior sequence of A-P states, starting in the most anterior EAD part of the axis and then going on to an A-P sequence of states in the head before ending in an A-P Hox sequence in the trunk and tail part of the axis (Fig 4).

**Fig 4.**
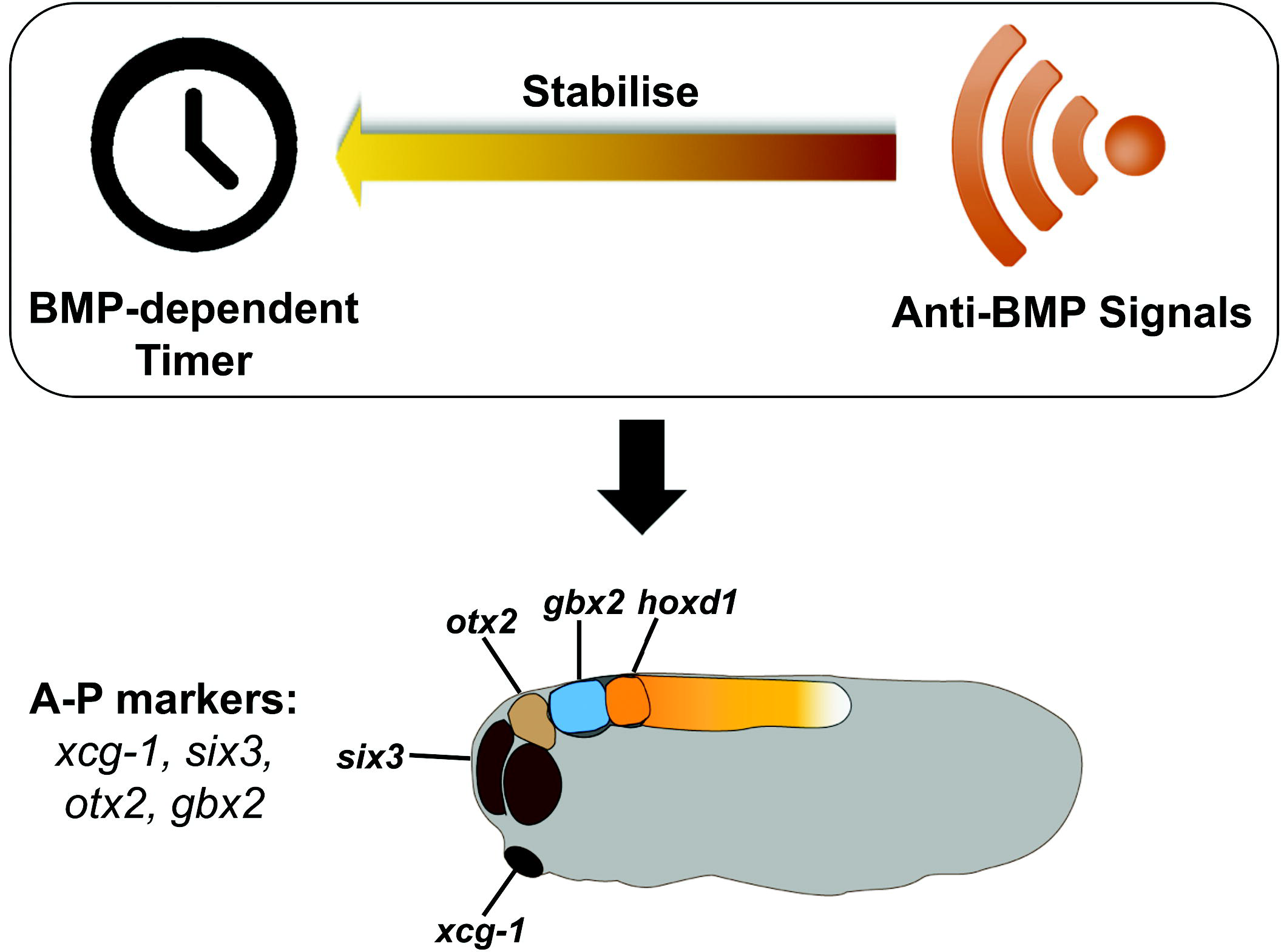
Hypothesis for head patterning by BMP signalling. A currently unknown, BMP-dependent timer is running during head patterning. The timer can be fixed by anti-BMP signals and regulate the expression of anterior genes: *xcg-1, six3, otx2, gbx2*, and *hoxd1*, resulting in the head being progressively patterned. Please note that the markers six3 and xcg1 are more ventrally placed than the rest. This is because the front end of the early A–P axis bends around ventrally to face backward like the handle of a walking stick (Durston 2015).

## Experimental procedures

### Microinjection

Frog embryos were harvested from naturally mated females and staged according to Nieuwkoop and Faber (Nieuwkoop & Faber 1994). For timed anti-BMP treatment in wild-type embryos, 200nL 0.1µg/µL human noggin protein (Sigma H6416) was injected into the blastocoel of embryos at stage 8, 9,10, 10.5 and 11. The embryos were then cultured to stage 28 for taking pictures. A similar approach has been used by others (Cooke & Smith 1989; Wacker *et al*. 2004a). mRNA for injection was transcribed with mMessage mMachine Kit (Ambion, Life technologies, AM1340) from the following plasmids after linearization at the appropriate restriction sites: pSP64T-BMP4 (for *BMP4* RNA) (Nishimatsu *et al*. 1992) and pCS2-hSmad6GR (for *smad6GR* RNA) (Marom *et al*. 2005). To induce full ventralisation, about 2ng *BMP4* RNA was injected to each embryo at 2-cell or 4-cell stage and cultured to stage 26. Timed anti-BMP treatment in BMP-ventralised embryos was achieved by combined injection of 2ng *BMP4* RNA and 2ng *smad6GR* RNA at 2-cell or 4-cell stage. The embryos were then treated with 10 µM dexamethasone for 2 hours at desired stages and cultured to stage 26.

### Whole mount in situ hybridization

When reached desired stages, embryos were fixed overnight in MEMFA at 4°C. After dehydration in 100% methanol, they were stored in methanol at −20°C until use. Whole mount in situ hybridization (WISH) was performed as previously described (Wacker et al., 2004a). The probes for in situ hybridization were synthesized from the following plasmids after linearization: pVZ1-xcg1 (for *xcg-1* probe) (Gammill & Sive 2000), pBSSK-Six3 (for *six3* probe) (Kenyon *et al*. 1999), pBluescript-KS-xotx2 (for *otx2* probe) (Blitz & Cho 1995), pXgbx-2 (for *gbx-2* probe) (von Bubnoff *et al*. 1996), and pBluescript SK-xHoxLab1 (for *hoxd1* probe) (Sive & Cheng 1991).

## Acknowledgements

We thank Dr A Fainsod for generously providing us with the *smad6GR* construct; Dr H Sive for the *xcg-1* probe and the *hoxd-1* probe; Drs S Moody, M Zuber, G Barsacchi and M Andreazzoli for the *six-3* probe; Dr K Cho for the *otx-2* probe; and Dr D Kimelman for the *gbx-2* probe.

