## Supplementary material for "Patterning of the vertebrate head in time and space by BMP signalling": Fig S1

**Supplementary figures**

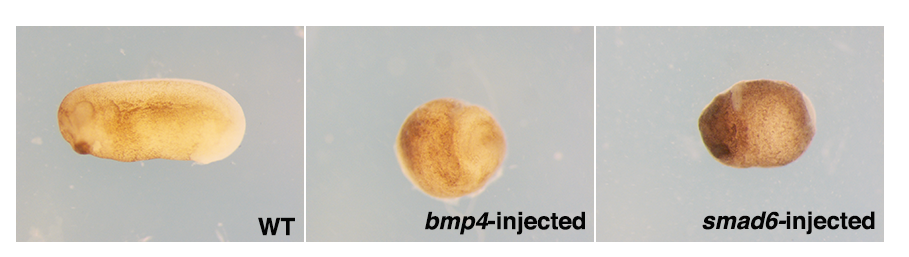

Figure S1: Ventralisation and dorsalisation by *bmp4* (high BMP) and smad6 (anti BMP) injection respectively. The *bmp4*-injected embryo formed a blob of tissue without axis. The smad6-injected embryo displayed a dorsalized phenotype, with cement gland forming at the front end of the embryo.
