## Supplementary material for "Patterning of the vertebrate head in time and space by BMP signalling": Fig S2

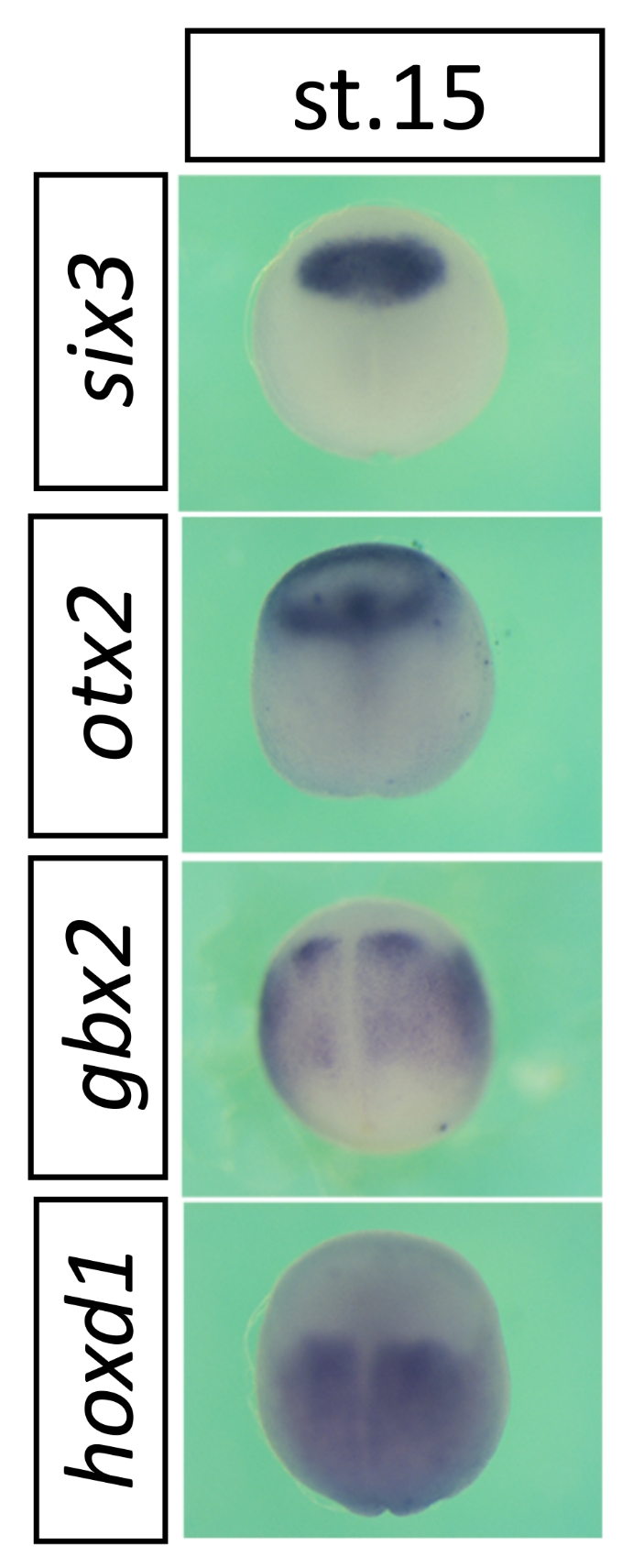

Figure S2: *six3*, *otx2*, *gbx2* and *hoxd1* are expressed sequentially from anterior to posterior. The expression of *six3*, *otx2*, *gbx2* and *hoxd1* in stage 15 embryos. Anterior is up.
